# Intracellular flow cytometry complements RT-qPCR detection of circulating SARS-CoV-2 variants of concern

**DOI:** 10.1101/2022.02.02.478775

**Authors:** Emiel Vanhulle, Becky Provinciael, Joren Stroobants, Anita Camps, Piet Maes, Kurt Vermeire

## Abstract

Despite the efficacy of current vaccines against SARS-CoV-2, the spread of the virus is still not under control, as evidenced by the ongoing circulation of the highly contagious SARS-CoV-2 Omicron variant. Basic and antiviral research on SARS-CoV-2 relies on cellular assays of virus replication *in vitro*. In addition, accurate detection of virus-infected cells and released virus particles is needed to study virus replication and to profile new candidate antiviral drugs. Here, by flow cytometry, we detect SARS-CoV-2 infection at single cell level and distinguish infected Vero E6 cells from uninfected bystander cells. Furthermore, based on the viral nucleocapsid expression, subpopulations of infected cells that are in an early or late phase of viral replication can be differentiated. Importantly, this flow cytometric technique complements RT-qPCR detection and can be applied to all current SARS-CoV-2 variants of concern, including the highly mutated Omicron variant.

**Method summary:** This study describes the characterization of SARS-CoV-2 infected cells using intracellular flow cytometric viral nucleocapsid staining that complements RT-qPCR quantification of viral RNA. The technique makes it possible to distinguish between infected cells in the early (low N) or late phase (high N) of viral replication. It can also be applied to the different variants of concern of SARS-CoV-2, including the Omicron variant.

## 1 Introduction

Since December 2019, Coronavirus Disease-2019 (COVID-19) has posed a serious threat to the international health and has impacted the global economy tremendously. The severe acute respiratory syndrome coronavirus 2 (SARS-CoV-2), a novel *Betacoronavirus*, was soon identified as the causative agent of COVID-19 (1, 2), and has quickly spread from the 41 initially reported patients in Hubei province, China to a global pandemic (3). As of February 1, 2022, there are nearly 373 million confirmed cases of SARS-CoV-2 infection and approximately 5.7 million deaths by COVID-19 worldwide (https://covid19.who.int/).

Despite the great success of the administered vaccines against SARS-CoV-2, the virus can still spread, as evidenced by the current circulation of the highly contagious Omicron variant (4). This emphasizes the additional need to develop effective antiviral countermeasures. Basic and antiviral research on SARS-CoV-2 relies not only on cellular assays for virus replication but also on the accurate detection of virus-infected cells and released virus particles. Here, we describe single cell quantification of SARS-CoV-2 infected cells by flow cytometry that complements the sensitive detection of viral RNA by RT-qPCR. We demonstrate the robustness of both techniques as they can detect various variants of SARS-CoV-2, including the (current circulating) highly mutated Omicron variant.

## 2 Materials and Methods

### 2.1 Cell lines and virus strains

#### Cell lines

African green monkey kidney cells (Vero E6 cells) were obtained from ATCC (CRL-1586) (Manassas, VA, USA) as mycoplasma-free stocks and were grown in Dulbecco’s Modified Eagle Medium (DMEM) (Thermo Fisher Scientific (TFS), Merelbeke, Belgium) supplemented with 10 % fetal bovine serum (FBS), 2 mM L-glutamine (TFS) and 0.075 % sodium-bicarbonate (TFS). Cells were maintained at 37 °C in a humidified environment with 5 % CO_2_, and were passaged every 3 to 4 days.

#### Viruses

All virus-related work was conducted in the high-containment biosafety level 3 facilities of the Rega Institute from the Katholieke Universiteit (KU) Leuven (Leuven, Belgium), according to institutional guidelines. Severe Acute Respiratory Syndrome coronavirus 2 isolates (SARS-CoV 2) were recovered from nasopharyngeal swabs of RT-qPCR-confirmed human cases obtained from the University Hospital Leuven (Leuven, Belgium). The following SARS-CoV-2 variants of concern were used in this study: Wuhan strain hCoV19/Belgium/GHB-03021/2020 (GISAID accession number EPI_ISL_407976), 20A.EU2 strain (clinical isolate A1), Alpha strain (UK-501Y.V1_B.1.1.7; EPI_ISL_791333), Beta strain (RSA-501Y.V2_B.1.351; EPI_ISL_896474), Gamma strain (BRA-501Y.V3_P.1; EPI_ISL_1091366), Delta strain (IND-B.1.617.2; EPI_ISL_2425097) and Omicron strain (RSA-B.1.1.529; EPI_ISL_7413964).

SARS-CoV-2 viral stocks were prepared by inoculation of confluent Vero E6 cells in a 150 cm^2^ culture flask at MOI of 0.02 in viral growth medium (DMEM supplemented with 2 % heat-inactivated FBS, 1 mM sodium Pyruvate and 1x MEM NEAA). After one hour incubation at 37 °C, viral growth media was added to bring the final volume to 25 mL. The cell supernatant was harvested at peak of infection, cleared by centrifugation and supernatant was aliquoted and stored at −80 °C.

To determine replication efficiency, the different SARS-CoV-2 strains were added to Vero E6 cells at an MOI of 0.02, as determined by end-point dilution titration on Vero E6 cells and calculated by the tissue culture infectious dose 50 (TCID50) method of Reed and Muench (5). The viral genome sequence was verified, and all infections were performed with passage 3 to 5.

### 2.2 Antibodies

The SARS-CoV-2 spike-specific monoclonal rabbit primary antibodies R001 and R007 were obtained from Sino Biological (Wayne, PA, USA; Cat. n° 40592-R001 and Cat. n° 40150-R007, respectively). The rabbit polyclonal SARS-CoV-2 nucleocapsid-specific antibody was obtained from GeneTex (Hsinchu City, Taiwan; Cat. n° GTX135357). The fluorescent-labelled Alexa Fluor 647 (AF647) goat anti-Rabbit IgG monoclonal antibody was from Cell Signaling Technologies (MA, USA; Cat. n° 4414).

### 2.3 Wild type virus infection assays

One day prior to the experiment, Vero E6 cells were seeded at 20,000 cells/well in 96 well-plates. Serial dilutions of the test compound were prepared in cell infectious media (DMEM + 2 % FCS), overlaid on cells and then virus was added to each well (MOI indicated in the figure legends). Cells were incubated at 37 °C under 5 % CO_2_ for the duration of the experiment. The supernatants were collected and stored at −20 °C.

### 2.4 RT-qPCR detection of SARS-CoV-2

Primers and probes for a duplex RT-qPCR were designed to detect the nucleocapsid (N) and envelope (E) gene of SARS-CoV-2 according to Centers for Disease Control and Prevention (CDC, USA; Cat. n° 2019-nCoVEUA-01) and Charité (Berlin, Germany), respectively (Table 1). Primers and probes were obtained from Integrated DNA Technologies (IDT, Leuven, Belgium). More specifically, for N2 the following probe was used: FAM-ACAATTTGCCCCCAGCGCTTCAG(BHQ1) and for E: HEX-ACACTAGCC(ZEN)ATCCTTACTGCGCTTCG(3IABkFQ). Final concentration of combined primer/probe mix consist of 500 nM forward and reverse primer and 250 nM probe. A stabilized *in vitro* transcribed universal synthetic single stranded RNA of 880 nucleotides in buffer with known copy number concentration (Joint Research Centre, European Commission, Cat. n° EURM-019) was used as a standard to quantitatively measure viral copy numbers.

**Table 1.**
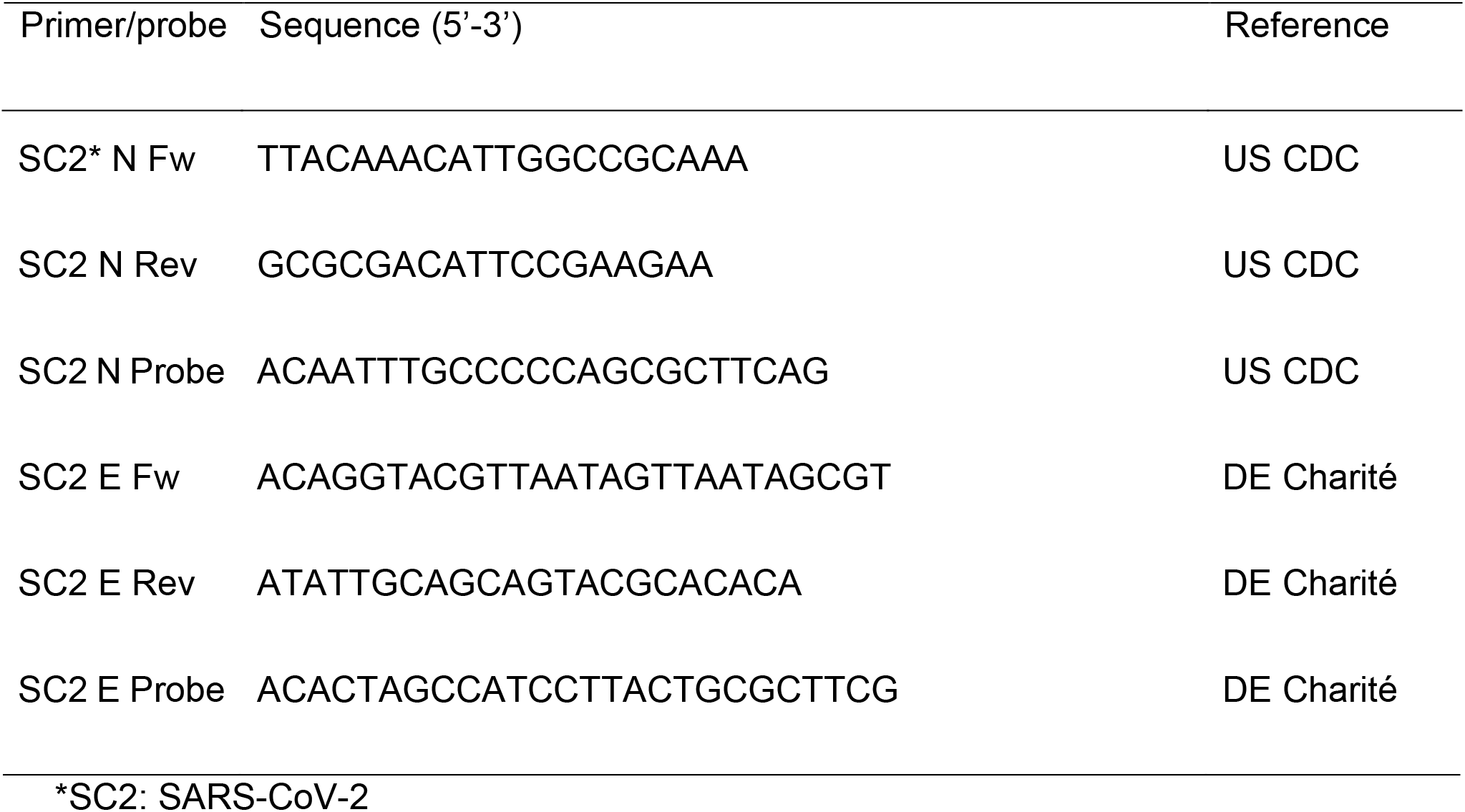
Primer and probe sequences used in duplex RT-qPCR

Total RNA was extracted from cell culture supernatants using QIAamp viral RNA mini kit (Qiagen) following manufacturer’s instruction. Viral E and N genes are simultaneously amplified and tested using a multiplex RT-qPCR. All the procedures follow the manufacturer’s instructions of the Applied Biosystems TaqMan Fast Virus one-step mastermix (ThermoFisher). Briefly, a 20 μl reaction mix was set up containing 5 μl of template, 7 μl of dH_2_O, 1.5 μl of each combined primer/probe mix (final concentrations of 500 and 250 nM, respectively) and 5 μl of 4X TaqMan Fast Virus 1-step mastermix (ThermoFisher). The qPCR plate was sealed and read in the FAM and HEX channels using a QuantStudio™ 5 Real-Time PCR system (ThermoFisher) under the following cycling protocol: 50°C for 5 min (reverse transcription), 95°C for 20 seconds (DNA polymerase activation), followed by 45 cycles of 95°C for 3 seconds (denaturation) and 55°C for 30 seconds (annealing and fluorescence collection) followed by an infinite 4°C hold.

### 2.5 Flow cytometry

For the intracellular staining of infected Vero E6 cells, cells were collected at different time points as indicated in the figure legends, and the Fixation/Permeabilization kit from BD Biosciences was used (Cat n° 554714). At the time of collection, supernatant was removed, and cells were washed in PBS. Then, trypsin (0.25 %) was added and plates were incubated for 3’ at 37 °C to detach the cells from the plate, followed by the addition of cold culture medium with 10 % FCS. Next, cells were resuspended, transferred to tubes and samples were centrifuged in a cooled centrifuge (4 °C) at 500 *g* for 5’. After removal of the supernatant, cells were fixed and permeabilized by the addition of 250 μL of BD Cytofix/Cytoperm buffer and incubated at 4 °C for 20’. Samples were washed twice with Perm/Wash buffer before the addition of the primary (anti-Nucleocapsid) antibody (0.3 μg per sample). After a 30 min incubation at 4 °C, samples were washed twice in BD Perm/Wash buffer, followed by a 30 min incubation at 4 °C with the secondary (labeled) antibody, and washed again. Finally, samples were stored in PBS containing 2 % formaldehyde (VWR Life Science AMRESCO). Acquisition of all samples was done on a BD FACSCelesta flow cytometer (BD Biosciences) with BD FACSDiva v8.0.1 software. Flow cytometric data were analyzed in FlowJo v10.1 (Tree Star). Subsequent analysis with appropriate cell gating was performed to exclude cell debris and doublet cells, in order to acquire data on living, single cells only.

## 3 Results and discussion

### 3.1 Optimization of duplex RTqPCR for SARS-CoV-2

At first, *in vitro* SARS-CoV-2 basic research in our laboratory was based solely on the cytopathic effect (CPE) readout of virus-infected Vero E6 cells (**Figure 1**). SARS-CoV-2 infection of a Vero E6 cell monolayer manifests in rounding and detaching of the cells. Administration of the neutralizing antibody R001, that binds to the receptor binding domain (RBD) of the viral spike (S) protein of SARS-CoV-2 (6), completely prevented the development of CPE, demonstrating a strong antiviral effect for R001 due to prevention of virus attachment to the cellular Angiotensin-Converting Enzyme 2 (ACE2) receptor (7) and subsequent virus entry in the cell. In contrast, the antibody R007, that also binds to the S protein but in a non-neutralizing manner, lacked antiviral potency (**Figure 1**).

**Figure 1.**
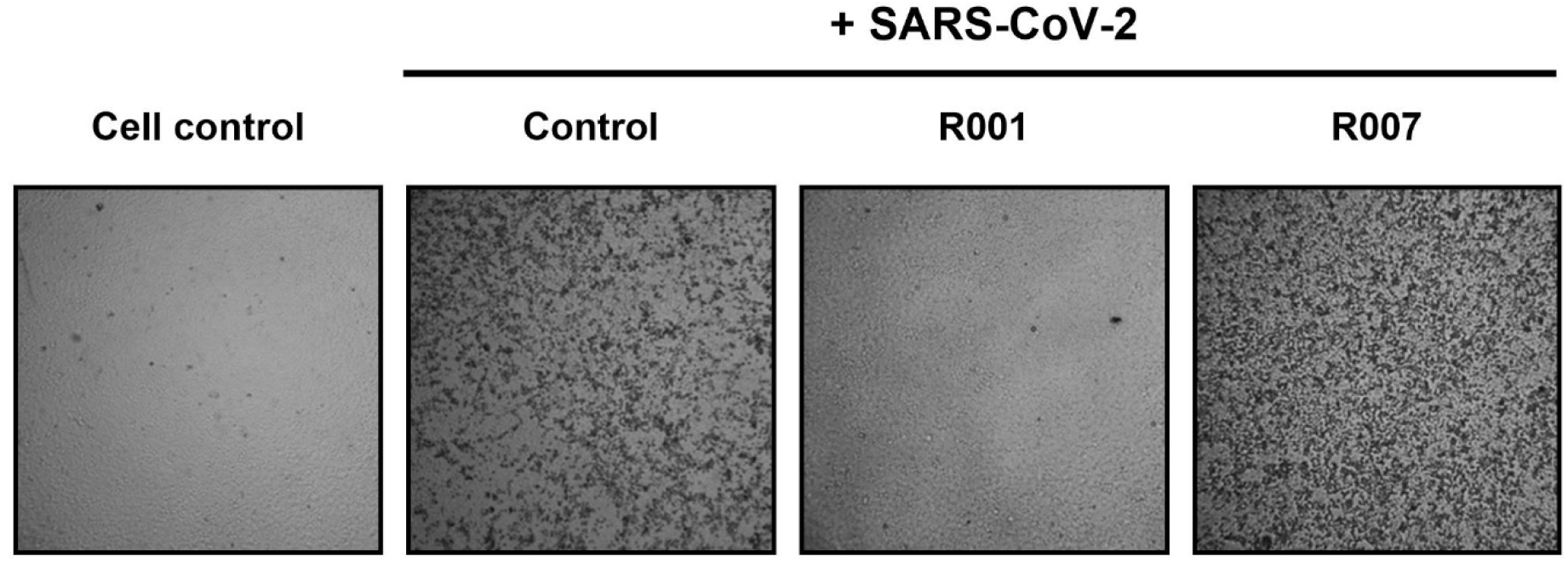
Cytopathic effect of SARS-CoV-2 virus on Vero E6 cells. Microscopic pictures of infected Vero E6 cells with SARS-CoV-2. Cells were exposed to SARS-CoV-2 (20A.EU2 variant; MOI = 0.015) for 2 hours, in the presence or absence of 2.5 μg/ml antibody R001 or R007. Supernatant was then removed, cells were washed and given fresh medium without (= ‘Control’) or with antibody (= ‘R001’ or ‘R007’). Pictures of cytopathic effect (CPE) were taken 3 days post infection. For R001 a complete protection of virus entry was obtained, whereas R007 had no antiviral effect. Magnification was 4X.

As CPE monitoring and scoring is a fairly subjective evaluation method, we next implemented a highly sensitive RT-qPCR method for the objective quantification of virus infection and virion production. To preserve the detection capacity of our assay for emerging SARS-CoV-2 mutants, we chose the simultaneous detection of two genes in the viral genome, i.e., the gene encoding for the highly abundant nucleoprotein N and that for the envelope protein E (**Figure 2A**). The primer sequences were selected to target conserved regions of the viral genome (for details see **Table 1**). Primer and probe sequences for N2 were obtained from Centers for Disease Control and Prevention (CDC, USA), whereas the sequences for E were from Charité (Berlin, Germany). In order to perform a duplex RT-qPCR, probes with matching fluorescent dyes (i.e., FAM and HEX) were designed accordingly. As proof of principle, we first validated our duplex RT-qPCR assay on a dilution series of a SARS-CoV-2 viral stock. Briefly, 1 over 10 dilutions of the virus stock in culture medium (starting from a 10^3^x dilution) were used for viral RNA extraction and subsequent analysis. The results of the E/N duplex RT-qPCR are shown in **Figure 2B**, demonstrating a similar accurate detection of virus samples with both genes in a concentration-dependent way (**Figure 2C**).

**Figure 2.**
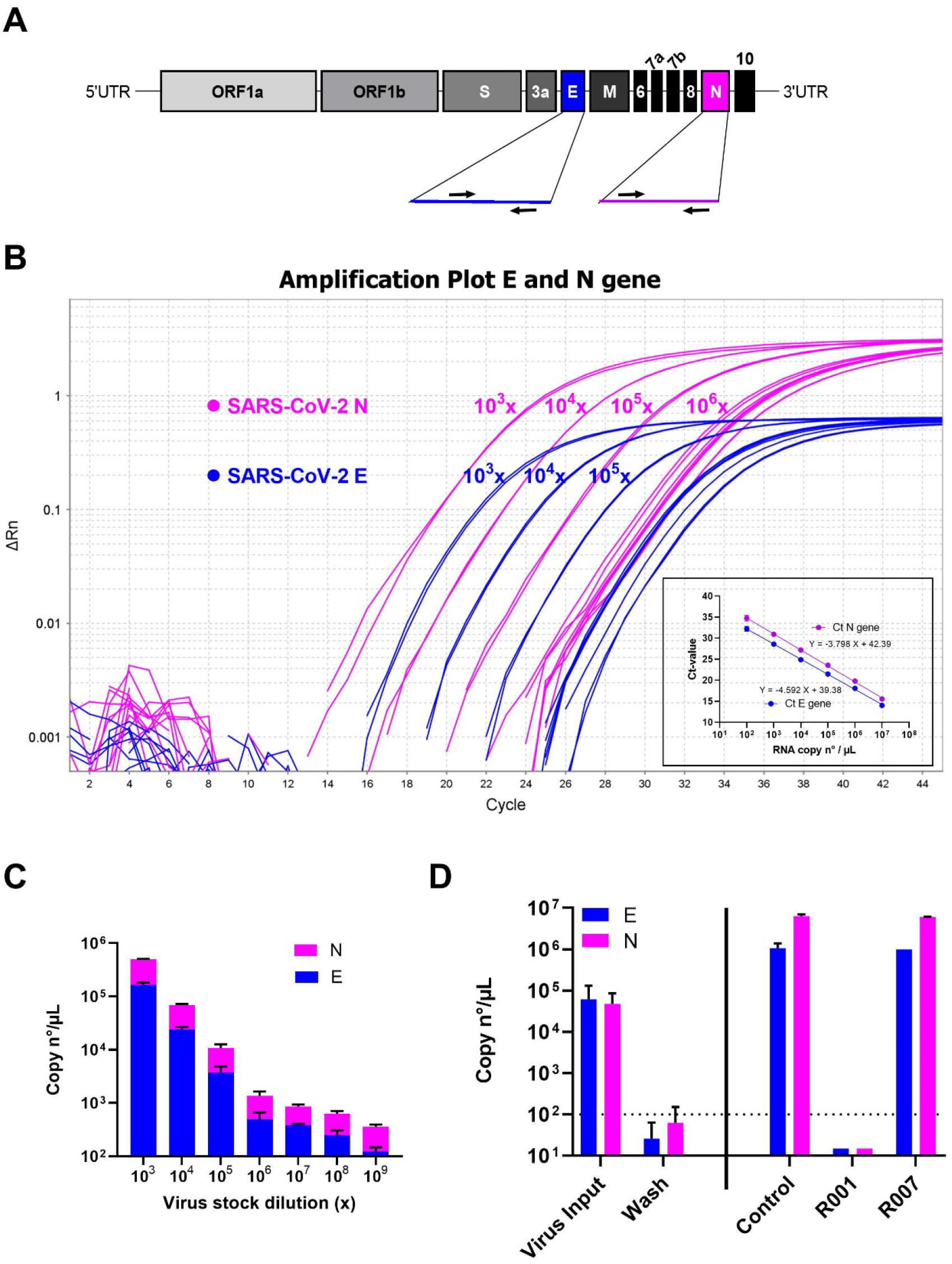
Duplex RT-qPCR detection of SARS-CoV-2 samples. **A)** Schematic representation of SARS-CoV-2 genome structure. The full-length RNA genome comprises approximately 30,000 nucleotides and has a replicase complex (comprised of ORF1a and ORF1b) at the 5’UTR. Four genes encode for the structural proteins, i.e., the Spike protein (S), the Envelope protein (E), the Membrane protein, and the Nucleocapsid protein (N). The genes for E and N were selected for RT-qPCR detection. **B)** Amplification plot of SARS-CoV-2 viral E and N gene. A dilution range (1:10) of a SARS-CoV-2 (20A.EU2 variant) virus stock was measured by duplex RT-qPCR for E and N gene (RNA) copy numbers, starting from a 10^3^x dilution. Graph shows amplification plot for 2 technical replicates of each sample. Insert shows the standard curve for the N and E gene RNA standards. **C)** Samples from (B) were analyzed for the quantification of copy numbers of E and N (mean ± SD, n = 2). **D)** RT-qPCR analysis of the samples from Figure 1. Vero E6 cells were exposed to SARS-CoV-2 (20A.EU2 variant; MOI = 0.015) for 2h. Supernatant (= virus input) was then removed, and cells were washed in PBS (= wash) and given fresh medium. In the conditions with the antibody-treatment (R001 or R007), 2.5 μg/ml of antibody was administered together with the virus, and administered again after the wash step. At day 3 post infection, supernatant was collected and a duplex RT-qPCR was performed to quantify the copy numbers of N and E. The dotted line refers to the residual amount (background) of virus that attached aspecifically to the cells. For R001 a complete protection of virus entry was obtained, whereas R007 had no antiviral effect. For each sample, the average (mean ± SD) of two technical replicates is given (n = 2), plotted on a log10 scale.

Next, we applied our duplex RT-qPCR analysis on the virus production of SARS-CoV-2 infected Vero E6 cells that were treated with virus-neutralizing antibodies. Briefly, virus yield analysis (i.e., new virions released by the infected cells in the culture medium) was performed at 3 days post infection (p.i.). Thus, supernatant of Vero E6 cells infected with SARS-CoV-2 was collected and subjected to RNA extraction and RT-qPCR. To assess if there was successful replication of the virus, we compared the samples with the virus input, i.e., the diluted virus stock solution to which the cells were exposed to for 2 hours. Also, we collected the wash sample (after the removal of virus input) that represents the amount of residual virus particles that were attached to the cells ‘aspecifically’ without productive viral entry in the cell, to determine the lower limit of detection of our virus yield assay. As shown in **Figure 2D**, Vero E6 cells were successfully infected with SARS-CoV-2 with productive virus release in the supernatant (‘Control’), as evidenced by the high copy numbers of viral RNA that exceeded the virus input. Treatment with antibody R001 completely abolished virus infection whereas R007 did not exert any antiviral activity, thus, confirming the CPE analysis of **Figure 1**. Thus, as comparable data were obtained for both SARS-CoV-2 genes, we concluded that we successfully developed a E/N gene duplex RT-qPCR.

To examine the robustness of our assay, we next determined the virus yield in Vero E6 cells infected with the different SARS-CoV-2 variants of concern. As summarized in **Figure 3**, our duplex RT-qPCR assay was able to simultaneously detect the viral genes E and N from all tested variants, also, from the current circulating Omicron variant. This also demonstrates that the many reported mutations in the different variants seem not to affect the primer binding sites of our assay.

**Figure 3.**
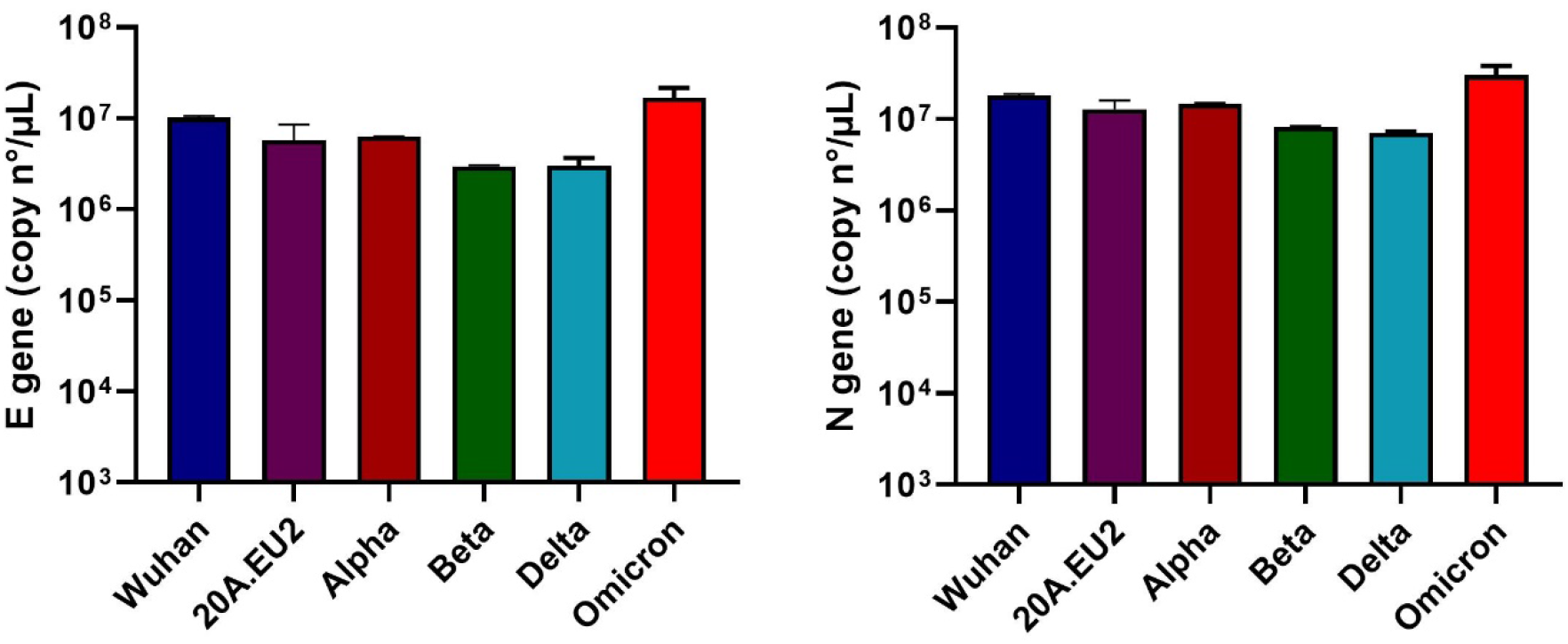
Duplex RT-qPCR analysis of virus yield from different SARS-CoV-2 variants of concern. Vero E6 cells were exposed to a fixed virus stock dilution (10x) of different variants of concern of SARS-CoV-2 (as indicated). After a 2 h incubation, supernatant (with virus) was then removed, and cells were washed in PBS and given fresh medium. At day 3 post infection, supernatant (that contained newly produced virus particles) was collected and a duplex RT-qPCR was performed to quantify the copy numbers of the E gene (left panel) and the N gene (right panel). For each condition, two biological replicates were included, and for the RT-qPCR quantification two technical replicates of each sample were analyzed to calculate the average value. Bars represent mean ± SD (n = 2).

### 3.2 Intracellular flow cytometric nucleoprotein staining of SARS-CoV-2 infected cells

Complementary to the RT-qPCR analysis described above, we implemented an assay to analyze virus infection at a single cell level, which is highly valuable to determine what cells from a heterogeneous cell population are susceptible to SARS-CoV-2. We opted for flow cytometry and selected an antibody that is raised against the viral nucleoprotein (8), a conserved viral protein that is abundantly expressed in virus-infected cells and stabilizes and protects the viral RNA (9).

Briefly, Vero E6 cells were infected with SARS-CoV-2, collected at selected time points p.i., fixed and permealized. Cells were then stained with a primary antibody against N, followed by incubation with a secondary antibody that is fluorescently labeled, and analyzed by flow cytometry. A kinetic study determined the optimal time point of cell collection, given that the cells need to be infected sufficiently, but not yet at that level of full blown CPE where the lytic effect of the virus destroys the infected cell population. As summarized in **Figure 4A**, viral N protein could be detected in Vero E6 cells at 24 hours p.i. (30 % infected cells), but longer incubation of the cells resulted in nearly a complete infected cell population (87 %). Interestingly, N detection by flow cytometry was not restricted to the SARS-CoV-2 Wuhan-related variant 20A.EU2, but also a 40 hours incubation of Vero E6 cells with the Gamma (Brazilian) variant returned a clear detection of viral N protein, showing an infection rate of 75 % (**Figure 4B**).

**Figure 4.**
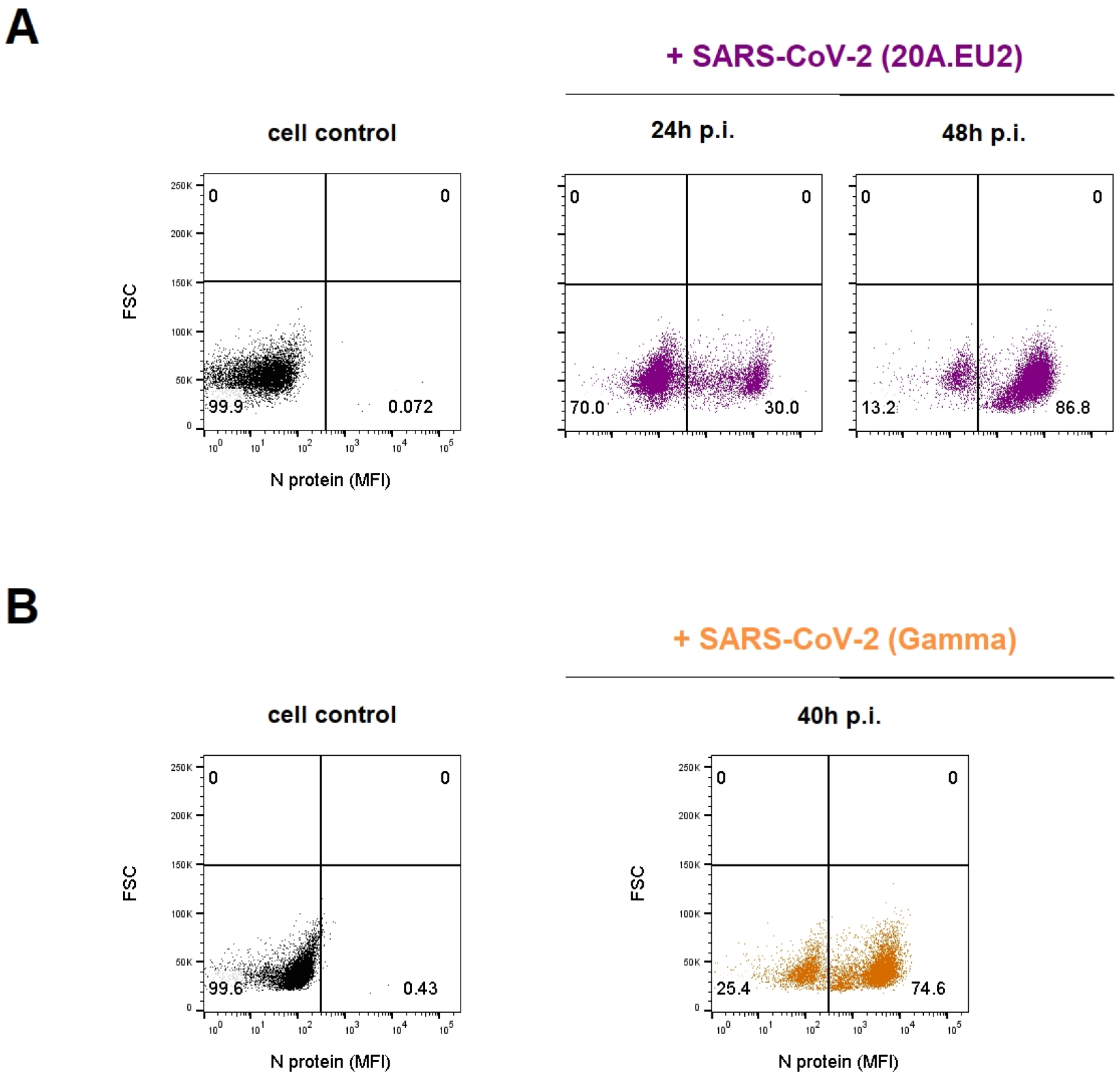
Kinetics of viral nucleoprotein expression in SARS-CoV-2 infected Vero E6 cells. **A)** Vero E6 cells were exposed to SARS-CoV-2 (20A.EU2 variant; MOI = 1.5) for 2h, washed and incubated with culture medium. At the indicated time points, cells were collected, fixed, permeabilized and stained with an antibody that binds to the viral nucleoprotein N. **B)** Same as in (A) but for the infection with the Gamma variant (MOI = 2). Dot plots show N expression in noninfected (lower left quadrant) and infected (lower right quadrant) Vero E6 cells, based on the analysis of 8,000 – 10,000 cells by flow cytometry. The numbers in each quadrant refer to the distribution of the cells (i.e., percentage of total cell population). The dot plot in black (left panel) represents the noninfected cell control.

Next, we assessed the robustness of this assay for the detection of different variants of concern of SARS-CoV-2 (**Figure 5**). For 6 different variants, including the recent circulating Omicron strain, a dilution range of virus stock was administered to Vero E6 cells. Next, 48 h p.i. cells were stained for viral N protein. As summarized in **Figure 5**, Vero E6 cells are highly permissive for all tested variants of concern of SARS-CoV-2, with infection efficiencies up to 68 %. For most variants with a concentrated virus stock (i.e., Wuhan, 20A.EU2, Beta and Omicron), dilution of virus input was related to higher infection rates. This discrepancy can be explained by the gradual increase in CPE with higher virus input which results in relatively more cell lysis and detached (infected) cells that are removed from the sample during wash steps. In most cases, the first dilution of the virus stock contains an excessive amount of virus that has a destructive impact on the cells. The longer survival of the infected cells in the condition of low virus input (right column of **Figure 5**) will be reflected by a higher relative number of infected cells as compared to the non-infected ones. Also, in this condition multiple rounds of virus infection can take place, resulting in a gradual increase in the absolute number of infected cells. For the Alpha variant, a 100x dilution seems to give the highest number of infected cells, whereas a more diluted virus input resulted in less infected cells. Of note, the virus stock of the Delta variant contained the least infectious particles, as evidenced by a concentration-dependent reduction in N positive cells. In fact, a 1,000x dilution of this stock returned an infection rate of less than 2 %. Importantly, our intracellular flow cytometry assay consistently detected all different variants of SARS-CoV-2, demonstrating the broad spectrum usage of this technique.

**Figure 5.**
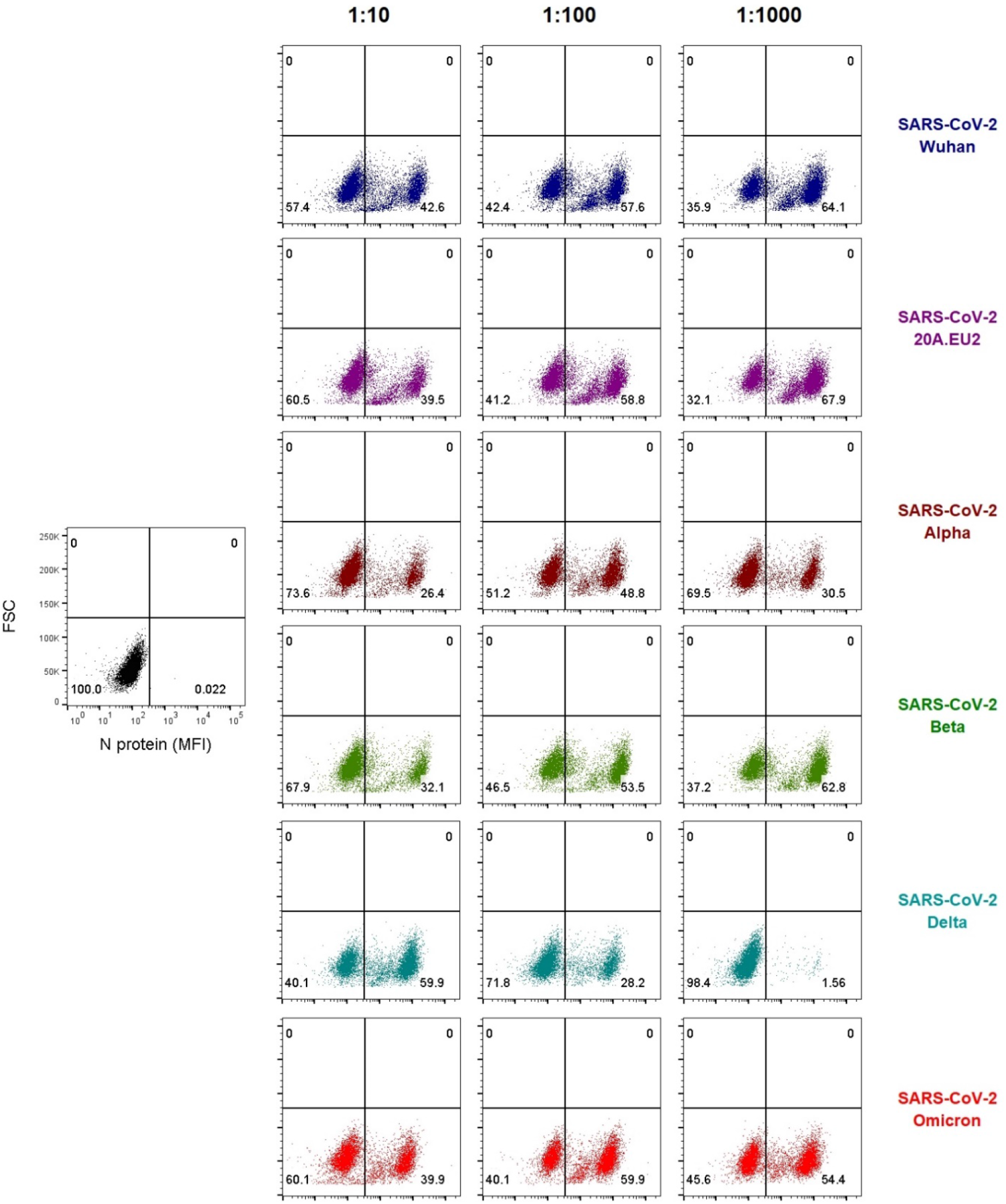
Viral N staining of Vero E6 cells infected with different variants of SARS-CoV-2. Vero E6 cells were exposed to a virus stock dilution (10x, 100x or 1000x) of different variants of concern of SARS-CoV-2 (as indicated). At day 2 post infection, cells were collected, fixed, permeabilized and stained with an antibody that binds to the viral N protein. Dot plots show N expression in noninfected (lower left quadrant) and infected (lower right quadrant) cells, based on the analysis of 8,000 – 10,000 Vero E6 cells by flow cytometry. The numbers in each quadrant refer to the distribution of the cells (i.e., percentage of total cell population). The dot plot in black (left panel) represents the noninfected cell control.

Finally, we analyzed the flow cytometric data in more detail. As shown in **Figure 6A**, infection of Vero E6 cells with low virus input of SARS-CoV-2 for 2 days resulted in a large population of infected cells. Moreover, two distinct cell populations of infected cells could be distinguished depending on the level of N protein expression. We hypothesized that the cells with low N protein levels are the newly infected cells in an early stage of the replication cycle. More specifically, those cells are infected cells that have taken up the virus but have not started the synthesis of new viral proteins yet. As the N protein is abundantly present in each virus particle to coat the viral genome (9), cells that contain virus particles should return detectable amounts of N protein. On the other hand, cells that are active replicating virus have started the biosynthesis of N protein and will produce N protein in high amounts (10). As the analysis of the flow cytometric data also includes a doublet discrimination gating step, the cells with high level N expression are not simple pairs of Vero E6 cells with double amount of viral N protein. To prove our hypothesis, and to investigate the nature of the two distinct N-positive cell populations, we performed an experiment in which we exposed Vero E6 cells to a high virus input (MOI of 28) for a short time (4 hours). This should allow enough time for the virus to enter the cells, but without starting the replication process (which is estimated at 6 to 8 hours post infection). As a control, we included the neutralizing antibody R001 as a virus entry inhibitor. As depicted in **Figure 6B**, already after 4 h p.i., a substantial amount (19 %) of virus-infected cells could be visualized (red square area on the dot plot), which was completely prevented by R001 treatment. Given that several wash steps were performed on the cells before the staining with the anti-N antibody, it is very unlikely that the N protein detected is from virus that has attached aspecifically to the cells without successful uptake. Furthermore, given the excess of virus, one should expect that all cells would then be ‘aspecifically’ coated with virus, resulting in a uniform population of infected cells. Of course, as Vero E6 cells are known for their efficient endocytic viral uptake mechanisms (11, 12), multiple viral particles can be taken up by a single cell. This can also explain the variation in N expression of the N low subpopulation. Longer incubation of the cells (24 h) resulted in the propagation of high N expressing cells (blue square area), thus, active virus-replicating cells. During this 24h time window, new virions will be released from the infected cells and can enter uninfected bystander cells, generating a new population of low N expressing cells. Thus, our flow cytometric analysis of SARS-CoV-2 infected cells can also indicate at a single cell level the status in the virus replication cycle. This information can be very helpful to profile new antiviral compounds as virus entry versus replication inhibitors.

**Figure 6.**
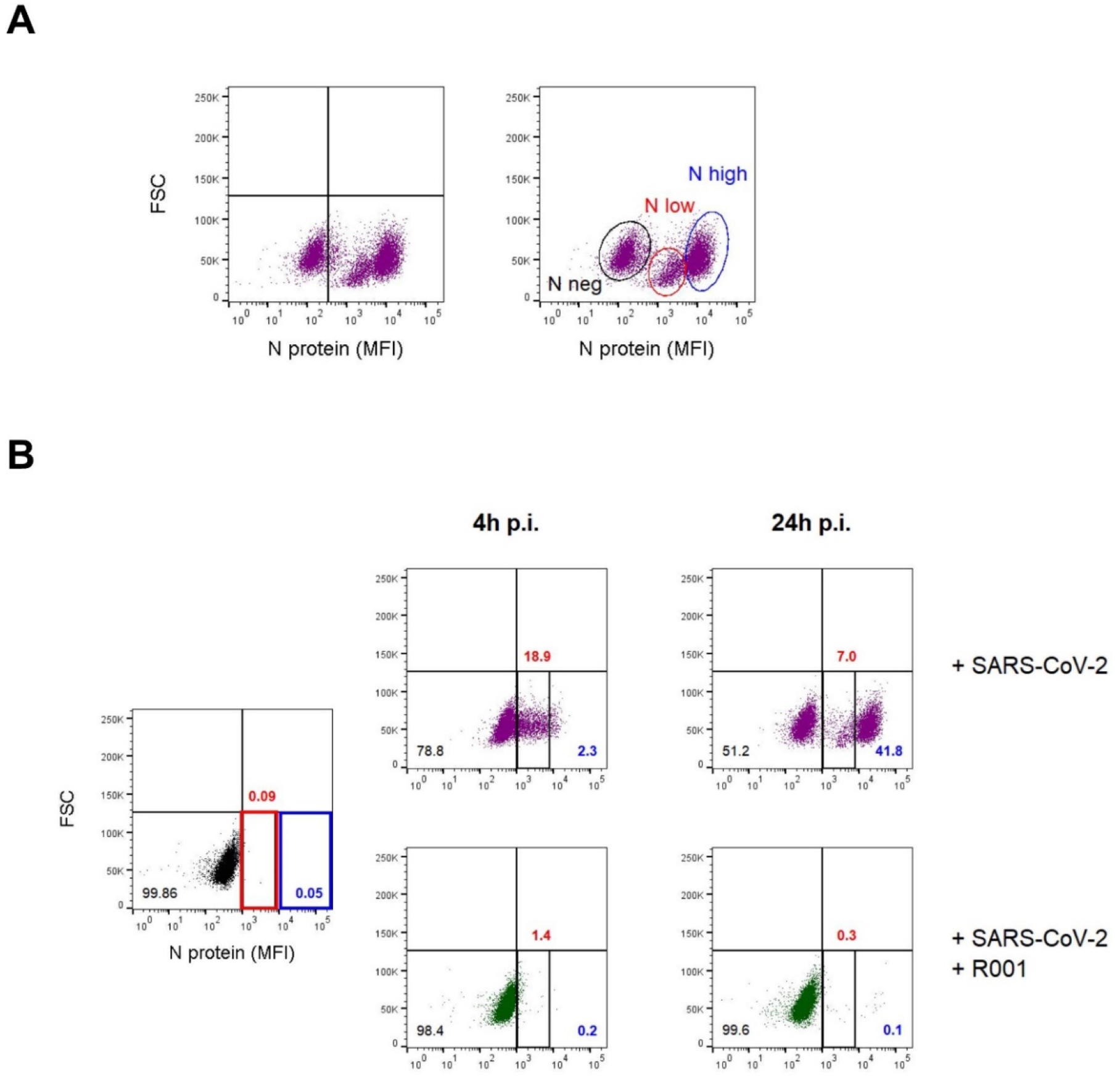
Viral N staining distinguishes two subpopulations of SARS-CoV-2 infected Vero E6 cells. **A)** Vero E6 cells were exposed to a low virus input of SARS-CoV-2 (20A.EU2 variant; MOI = 0.28). At day 2 post infection, cells were collected, fixed, permeabilized and stained with an antibody that binds to the viral nucleoprotein N. Dot plots show N expression in noninfected (lower left quadrant) and infected (lower right quadrant) cells, based on the analysis of 8,000 – 10,000 Vero E6 cells by flow cytometry. The graph shows two distinct subpopulations of infected cells, i.e., “N low” (marked by a red line) and “N high” (marked by a blue line). **B)** Vero E6 cells were exposed to a high virus input of SARS-CoV-2 (20A.EU2 variant 1:10; MOI = 28), without (top panels) or with 10 μg/ml of neutralizing antibody R001 (bottom panels). At 4h post infection, half of the samples were washed to remove virus input and were administered fresh medium (without or with R001) and incubated for another 20h followed by N staining. The other half of the cells were collected at 4h p.i., fixed, permeabilized and stained with an antibody that binds to the viral N protein. Dot plots show N expression in infected cells, based on the analysis of 8,000 – 10,000 Vero E6 cells by flow cytometry. The two distinct subpopulations of infected cells, i.e., “N low” (marked in red) and “N high” (marked in blue) are indicated. The numbers refer to the distribution of the cells (i.e., percentage of total cell population). The dot plot in black (left panel) represents the noninfected cell control.

## Conclusions and future perspective

In the context of basic research on viruses, a repertory of cell-based assays and sensitive techniques for virus detection is requested. In this study, we successfully optimized a SARS-CoV-2 duplex RT-qPCR for the detection of viral RNA copies by the simultaneous amplification of the nucleocapsid and envelope genes. This sensitive technique was able to detect viral E and N from all tested variants, also, from the current circulating Omicron variant. This demonstrates that the many reported mutations in the different variants seem not to affect the primer/probe binding sites of our PCR. For the quantification of virus replication and virion production, samples for RT-qPCR are mostly based on cell culture supernatant from infected cells or cell lysate, thus originating from thousands of cells, either infected or uninfected. However, with our flow cytometric technique, based on intracellular viral nucleocapsid staining, we succeeded to analyze virus infection at the single cell level, which is highly valuable to determine what cells from a heterogeneous cell population are susceptible to SARS-CoV-2. Furthermore, this technique makes it possible to distinguish between subpopulations of infected cells that are in an early (low N) or late phase (high N) of viral replication. This can be an added value in the profiling of new candidate antiviral drugs, to discriminate between entry inhibitors and viral polymerase inhibitors that block viral replication. Importantly, this flow cytometric technique complements RT-qPCR detection and can be applied to all current SARS-CoV-2 variants of concern, including the highly mutated Omicron variant. We believe that this method will be a valuable supplement for the ongoing study of COVID-19.

## Author contributions

K.V., E.V. and J.S. designed research; E.V., A.C., B.P. and J.S. performed research; E.V. and K.V. analyzed the data; E.V. and K.V. wrote the manuscript; P.M. contributed new reagents/analytic tools and critically reviewed the manuscript. All of the authors discussed the results and commented on the manuscript.

## Acknowledgements

We thank Geert Schoofs for technical support to the flow cytometry experiments and Dominique Schols for financial support.

## Information pertaining to writing assistance

N/A

## Ethical disclosure

The authors state that they have obtained appropriate institutional review board approval or have followed the principles outlined in the Declaration of Helsinki for all human or animal experimental investigations.

## Data sharing statement

N/A

